# Harnessing the power of eDNA metabarcoding for the detection of deep-sea fishes

**DOI:** 10.1101/2020.07.10.197012

**Authors:** Beverly McClenaghan, Nicole Fahner, David Cote, Julek Chawarski, Avery McCarthy, Hoda Rajabi, Greg Singer, Mehrdad Hajibabaei

**Author notes:** Corresponding author (MH).

## Abstract

The deep ocean is the largest biome on Earth and faces increasing anthropogenic pressures from climate change and commercial fisheries. Our ability to sustainably manage this expansive habitat is impeded by our poor understanding of its inhabitants and by the difficulties in surveying and monitoring these areas. Environmental DNA (eDNA) metabarcoding has great potential to improve our understanding of this region and to facilitate monitoring across a broad range of taxa. Here, we evaluate two eDNA sampling protocols and seven primer sets for elucidating fish diversity from deep sea water samples. We found that deep sea water samples (> 1400 m depth) had significantly lower DNA concentrations than surface or mid-depth samples necessitating a refined protocol with a larger sampling volume. We recovered significantly more DNA in large volume water samples (1.5 L) filtered at sea compared to small volume samples (250 mL) held for lab filtration. Furthermore, the number of unique sequences (exact sequence variants; ESVs) recovered per sample was higher in large volume samples. Since the number of ESVs recovered from large volume samples was less variable and consistently high, we recommend the larger volumes when sampling water from the deep ocean. We also identified three primer sets which detected the most fish taxa but recommend using multiple markers due the variability in detection probabilities and taxonomic resolution among fishes for each primer set. Overall, fish diversity results obtained from metabarcoding were comparable to conventional survey methods. While eDNA sampling and processing need be optimized for this unique environment, the results of this study demonstrate that eDNA metabarcoding can be employed to facilitate biodiversity surveys in the deep ocean, require less dedicated survey effort per unit identification and are capable of simultaneously providing valuable information on other taxonomic groups.

## Introduction

The deep ocean is the largest biome on Earth by volume and also one of the planet’s most understudied environments [1]. The biodiversity of the deep ocean has not been fully explored nor is the distribution and biology of many deep-water species well understood [1–3]. Despite our limited knowledge of deep-water fauna, several species are commercially targeted and, along with many other taxa, face increasing pressure from climate change [4–6]. Monitoring and managing the impacts of commercial fishing and climate change in this environment is difficult due to logistic constraints and the high cost of sampling such challenging environments [7]. Despite these impediments, documenting the biodiversity of this region is integral to sustainable management and ecosystem monitoring.

Deep ocean biodiversity surveys are often done using a combination of methods, each targeting a particular taxonomic group. For fish and micronekton, trawling, long-lining, and acoustic monitoring are often used. Small nets and filtration systems can target small zooplankton and phytoplankton and autonomous video camera systems can capture a range of macrofauna [8]. Each of these methods have limitations in their ability to capture a community based on morphological and behavioral selectivity as well as taxonomic resolution. Additionally, not all of these methods can be employed equally well in all areas of the ocean. For example, bottom trawling is ineffective for surveying along steep slopes and rocky surfaces and is undesirable in areas with sensitive epifauna, such as deep-water corals and sponges [9,10]. The need to employ multiple sampling methods to assess the biodiversity of the deep sea increases the sampling effort required, complicates the interpretation of data, and thereby adds to the challenges of surveying this environment.

Metabarcoding using environmental DNA (eDNA) is a relatively new approach to biodiversity analysis that can facilitate surveys by reducing the sampling effort and taxonomic expertise required and thus far metabarcoding has been underutilized in the deep ocean [11]. Marine eDNA studies have primarily surveyed coastal and/or surface water (e.g. [12,13]), with very few studies sampling water at depths > 1000m for eukaryotic eDNA (but see [14]). Much of the eDNA work on deep-sea communities has focused on sediment sampling to study benthic communities (e.g. [15–17]) as opposed to fish and pelagic communities.

Using eDNA from deep sea water samples to characterize biodiversity has the potential to provide critical insight into deep ocean biodiversity however, eDNA sampling protocols need to be optimized for this environment. Abiotic factors, such as the reduced light levels and comparatively low variability in temperature and salinity in deep ocean water [2], affect the persistence of eDNA while biotic factors, such as the predominant life histories and/or metabolism of the organisms living in the deep ocean (e.g. slower metabolism; [18]), may affect the amount of eDNA released into the water. Therefore, the optimal protocols for eDNA sampling in the deep ocean must be determined separately from coastal and surface marine water sampling and furthermore, sample processing should be optimized for the particular target groups of deep-sea organisms (e.g. fish).

The objectives of this study were to develop an eDNA metabarcoding sampling protocol for the deep sea, evaluate the performance of multiple primer sets for the detection of deep-sea fishes and compare eDNA results to conventional fish surveys. We collected seawater samples over two sampling seasons and refined the sampling and lab protocols in the second season to improve the detection of deep-sea fishes. While fishes were the target group, we also report general biodiversity results that were detected concurrently.

## Methods

### Study Area

We surveyed fish communities in the Labrador Sea, in the Northwest Atlantic Ocean, in the summer (June-August) over three sampling years. Surveys using conventional sampling techniques (2017-2019) and eDNA water sampling (2018-2019) were conducted along three transects each covering a water depth gradient of approximately 500 m to 3000 m (see S1 Fig for map and S1 Table for GPS coordinates). All field sampling was conducted under experimental licenses from Fisheries and Oceans Canada.

### Conventional Fish Surveys

Harvester logbooks and research vessel (RV) surveys using Campelen trawls are typically used to monitor and manage demersal fish communities in Canadian waters but these collections are restricted to waters less than 1500m and are relatively sparse for northern areas [19]. In deeper waters (>1500 m) of the Labrador Sea, there is very limited information on demersal or pelagic fish communities. Therefore, to augment species lists from RV surveys and logbooks, targeted sampling of demersal (baited hooks and cameras; [19]) and pelagic (Isaac Kidd Midwater Trawls (IKMT) [20]) fish was conducted in the study area. Demersal fish sampling was conducted along two transect lines in 2017 and 2019, whereas pelagic fish communities were sampled across three transect lines in 2018 and 2019. Baited hooks and cameras were deployed on the ocean bottom whereas IKMT samples were collected from the mesopelagic deep-scattering layer (an area of concentrated pelagic biomass [21]; sampled depths ranged from 360 – 536 m) as detected by hull-mounted echosounders. Fish captured using both methods were identified morphologically. While the exact sampling sites differed for pelagic and demersal sampling sites, pelagic sampling was conducted over the same transects as the demersal sampling but was restricted to a maximum water depth of 2500 m (versus a maximum depth of ~3000 m for demersal sampling).

### eDNA Water Sample Collection

eDNA water samples were collected from seven stations along one transect in 2018. In 2019, two of these stations were resampled and water samples were collected from an additional eight stations along the two other transects. At each station, samples were collected from the surface, the deep scattering layer and just above the bottom up to a depth of ~2,500 m depth (n = 144, S1 Table). Water samples were co-located in time and space with pelagic fish (IKMT) sampling. Samples were collected using a Niskin-style rosette sampler. Rosette bottles were assigned to eDNA sampling for the duration of the field mission and were decontaminated prior to sampling and between stations using ELIMINase (Decon Labs). At each sampling station, a field blank was collected using distilled water to control for potential contamination. In 2018, we employed a sampling strategy adapted from previous coastal surface water sampling in the North Atlantic [22], where triplicate 250 mL samples were collected at each sampling depth. Water samples were then frozen at −20°C and shipped frozen to the lab for subsequent processing. Water filtration took place in a clean lab, thereby reducing the potential for sample contamination, however cold storage space was required on the vessel to store water samples. Based on the results of 2018 sampling, the sampling strategy was modified for 2019. We increased the water volume collected by a factor of 6, collecting triplicate 1.5 L water samples in 2019, however the larger volume of water collected could not be kept in cold storage on the vessel due to space limitations. As such, water samples in 2019 were filtered on the vessel. Filter cartridges (requiring less storage space) were stored at −20°C for the duration of the expedition.

### Laboratory Procedures

All water samples were filtered through 0.22 μm PVDF Sterivex filters (MilliporeSigma) using a peristaltic pump. Filtration on the vessel took place in a dedicated lab space that included a positive pressure ventilation system. Before each filtration session, surfaces and equipment were all decontaminated with ELIMINase and rinsed with deionized water. Filtration began immediately after sample collection (average volume filtered 1.35 ± 0.15 L). For samples filtered in the lab, filtration took place in a PCR clean lab under a laminar flow hood (AirClean Systems) which was decontaminated using ELIMINase, lab-grade water and 70% ethanol prior to each sample set. Water samples were thawed at 4⍰°C and immediately filtered. DNA was extracted from all filter membranes using the DNeasy PowerWater Kit (Qiagen). DNA extracts were quantified using the Quant-iT PicoGreen dsDNA assay with a Synergy HTX plate fluorometer (BioTek).

Seven DNA markers from three gene regions (cytochrome c oxidase I (COI), 12S and 18S) were selected to assess eukaryotic biodiversity in the 2018 samples (Table 1A), including three primer sets specifically for bony fish. The 2019 samples were analyzed with only these three fish-targeting primer sets. Each PCR reaction contained 1X reaction buffer, 2 mM MgCl_2_, 0.2mM dNTPs, 0.2 μM of each of the forward and reverse Illumina-tailed primers, 1.5U Platinum Taq (Invitrogen) and 1.2 μL of DNA in a total volume of 15 μL. Due to the higher concentration of DNA recovered from 2019 samples, diluted DNA was used for 2019 samples (1/10 and 1/2 for surface samples and samples at depth, respectively). See Table 1B for PCR conditions for all primer sets. Three PCR replicates were performed for each primer set from each sample and then pooled for a single PCR cleanup with the QIAquick 96 PCR purification kit (Qiagen).

**Table 1.**
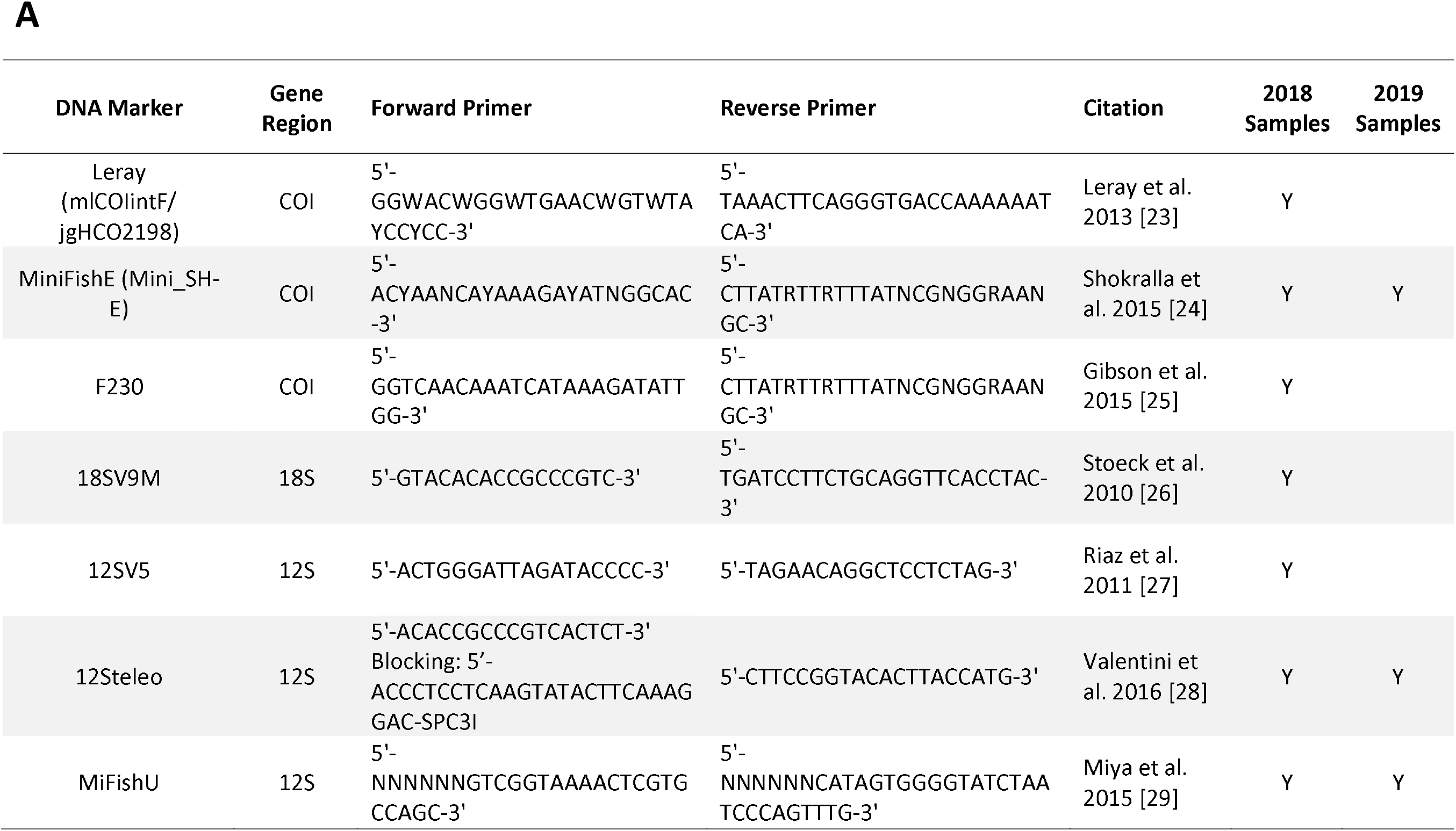
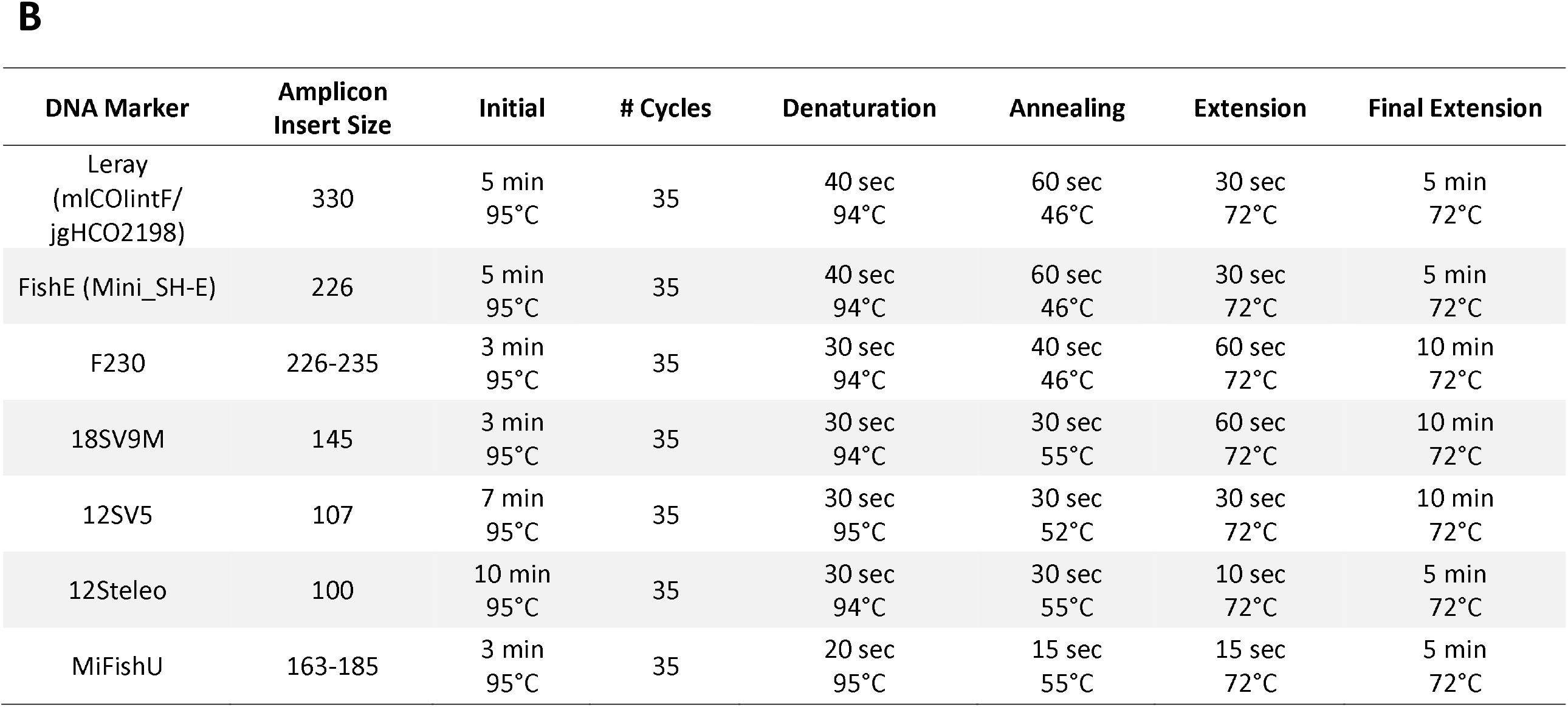
Marker summary indicating (A) the gene region, primer sequences, citation, and sample set(s) (2018 and 2019) the marker was used on and (B) the amplicon insert size and PCR conditions used for each marker.

**Table 1.** ESV level summary of taxonomic identifications via metabarcoding for each primer set. # ESV indicates the total number of ESVs detected, # ESV Tax represents number of ESVs with taxonomic matches at any level, # Metazoan ESV indicates the number of ESVs identified as Metazoa (≥0.9 selection criteria, kingdom = Metazoa) and # Fish ESV indicates the number of ESVs identified as fish (> 0.9 selection criteria, class = Actinopteri or Chondrichthyes). Table (A) summarizes the number and percentage of metazoan ESVs assigned to each taxonomic level and table (B) summarizes the number and percentage of fish ESVs assigned to each taxonomic level.

| Primer Set | # ESV | # ESV Tax | # Metazoan ESV | Family |  | Genus |  | Species |  |
| --- | --- | --- | --- | --- | --- | --- | --- | --- | --- |
|  |  |  |  | # ESV | % | # ESV | % | # ESV | % |
| 12S | 890 | 245 | 70 | 69 | 98.6% | 68 | 97.1% | 14 | 20.0% |
| 12Steleo | 1,192 | 1,106 | 312 | 309 | 99.0% | 184 | 59.0% | 22 | 7.1% |
| 12S MiFishU | 13,228 | 12,935 | 73 | 69 | 94.5% | 58 | 79.5% | 14 | 19.2% |
| 18SV9M | 7,081 | 6,878 | 72 | 38 | 52.8% | 24 | 33.3% | 7 | 9.7% |
| COI F230 | 16,252 | 15,952 | 57 | 0 | 0.0% | 0 | 0.0% | 0 | 0.0% |
| COI FishE | 89,506 | 61,647 | 498 | 422 | 84.7% | 166 | 33.3% | 70 | 14.1% |
| COI Leray | 20,190 | 16,628 | 44 | 40 | 90.9% | 35 | 79.5% | 22 | 50.0% |

**B**
| Primer Set | # ESV | # ESV Tax | # Fish ESV | Family |  | Genus |  | Species |  |
| --- | --- | --- | --- | --- | --- | --- | --- | --- | --- |
|  |  |  |  | # ESV | % | # ESV | % | # ESV | % |
| 12S | 890 | 245 | 2 | 2 | 100.0% | 2 | 100.0% | 1 | 50.0% |
| 12Steleo | 1,192 | 1,106 | 110 | 107 | 97.3% | 49 | 44.5% | 7 | 6.4% |
| 12S MiFishU | 13,228 | 12,935 | 29 | 27 | 93.1% | 21 | 72.4% | 4 | 13.8% |
| 18SV9M | 7,081 | 6,878 | 9 | 2 | 22.2% | 0 | 0.0% | 0 | 0.0% |
| COI F230 | 16,252 | 15,952 | 2 | 0 | 0.0% | 0 | 0.0% | 0 | 0.0% |
| COI FishE | 89,506 | 61,647 | 19 | 16 | 84.2% | 12 | 63.2% | 9 | 47.4% |
| COI Leray | 20,190 | 16,628 | 0 | 0 | 0.0% | 0 | 0.0% | 0 | 0.0% |

Amplicons were visualized using agarose gel (1.5% w/v) electrophoresis to verify amplification of DNA markers and to assess negative controls generated during PCR, extraction, filtration, and field collection. Negative controls were carried through to sequencing as an added level of verification. Amplicons were then indexed using unique dual Nextera indexes (IDT; 8-bp index codes). Indexing PCR conditions were initiated for 3 mins at 95⍰°C, followed by 12 cycles of 95⍰°C for 30⍰s, 55⍰°C for 30⍰s, and 72⍰°C for 30⍰s, and a final extension at 72⍰°C for 5⍰mins. Amplicons were quantified with Quant-iT PicoGreen dsDNA assay and pooled together in equimolar concentrations by DNA marker. Amplicon pools were cleaned using AMPure XP cleanups, quantified with a Qubit fluorometer (Thermo Fisher) and the size distribution of each pool was verified with the DNA 7500 kit on the Agilent 2100 Bioanalyzer. The 2018 12SV5, COI Leray, COI MiniFishE, 18SV9M, and COI F230 amplicon pools were combined into one library. The 2019 12Steleo, 12S MiFishU and COI MiniFishE amplicons pools were combined with the 2018 12Steleo and 12S MiFishU amplicon pools in a second library. The libraries were sequenced with a 300-cycle S1 kit and a 500-cycle SP kit, respectively, on the NovaSeq 6000 following the NovaSeq standard workflow with a target minimum sequencing depth of 1 million sequences per sample per amplicon. Raw sequence reads are available in NCBI’s sequence read archive under project PRJNA643526.

### Bioinformatics

Base calling and demultiplexing were performed using Illumina’s bcl2fastq software (v2.20.0.422). Primers were trimmed from sequences using *cutadapt* v1.16 [30] and then DADA2 v1.8.015 [31] was used for quality filtering, joining paired end reads (maxEE⍰=⍰2, minQ⍰=⍰2, truncQ⍰=⍰2, maxN⍰=⍰0) and denoising using default parameters to produce exact sequence variants (ESVs). Taxonomy was assigned to ESVs using NCBI’s blastn tool v1.9.0 [32] and the nt database (downloaded: November 30, 2019) with an e-value cut-off of 0.001. In cases where a sequence matched multiple taxa with an equally high score, we only assigned taxonomy to the lowest common ancestor of the ambiguous hits. The resulting taxonomic hits were filtered using a selection criterion (% sequence similarity x length of overlap). Family-level matches were reported using a minimum of 95% selection criterion, genus-level matches were reported using a minimum of 98% selection criterion and species-level matches were reported using a 100% or perfect match. All taxa detected were verified using the WoRMS [33] and EOL [34] databases and spurious or irrelevant hits (e.g. terrestrial or domestic species) were omitted.

### Statistical Analysis

All statistical analyses were performed using R v3.5.1 [35]. Sampling sites in 2018 (250 mL) and 2019 (1.5 L) did not overlap completely therefore samples with different volumes were collected in different locations and different years, meaning no direct comparisons between samples can be made. However, we made general comparisons across all small volume samples and all large volume samples. Additionally, previous studies in this region suggest that spatial differences in community structure are small compared to community changes by water depth [19]. We used a robust two-way ANOVA (α = 0.05) implemented using the *‘Rfit’* package v0.24.2 [36] to compare the DNA concentrations in each sample between sampling volumes and between sampling depths, categorized as shallow (<500 m), mid-depth (500-1400 m) or deep (>1400 m). Shallow includes samples from the surface and the deep scattering layer for some stations. Mid-depth includes samples from the deep scattering layer and the bottom for some stations. Deep includes only samples from the bottom. Depth categories were chosen based on the distribution of depths sampled at each site and preliminary data exploration (see S2 Fig). Using data from the three markers used on 2018 and 2019 samples (COI MiniFishE, 12Steleo, 12S MiFishU), we used a robust two-way ANOVA to compare the number of ESVs recovered in each sample between sampling volumes and between sampling depths. Post-hoc comparisons between groups were performed using the *‘rcompanion’* package v1.13.2 [37]. We used Levene’s test to determine if the variance in DNA concentration and number of ESVs differed between years and water depths.

We assessed the performance of different primer sets by comparing the number of taxa detected and the resolution of taxonomic assignments for all markers, with a particular focus on the recovery and resolution of fishes. In addition, we used a multi-species, multi-scale occupancy modeling framework to compare the detection probabilities of all fish specific primer sets (12S MiFishU, 12Steleo, COI MiniFishE) across fish taxa while accounting for false negatives following McClenaghan et al. [38]. We included water depth (meters) as a covariate at the level of occupancy and primer set as covariate at the level of detection probability (see S1 File for model formulation and detailed methods). We ran two models using observations from different levels of taxonomic resolution: fish species and fish families.

We compared the fish taxa detected via eDNA metabarcoding to the fish taxa detected via conventional survey methods for a single sampling expedition (2019 eDNA and 2019 pelagic IKMT sampling). This represents equal field sampling effort for both methods. We summarized the total number of taxa detected using each method at multiple taxonomic levels (family, genus, and species). Additionally, we summarize the total number of taxa recovered from multiple years and methods of conventional sampling.

## Results

### General Sequencing Summary

The mean number of sequences recovered per sample per amplicon after bioinformatic filtering was 1,250,418 (range: 16 – 13,603,412) yielding an average of 706 (range: 1-6003) ESVs per sample per amplicon and a total of 148,339 ESVs. 77.8% of the ESVs matched a sequence in the reference database, although the taxonomic rank assigned to each ESV was variable and resolution differed between amplicons (Table 2).

**Table 2.**
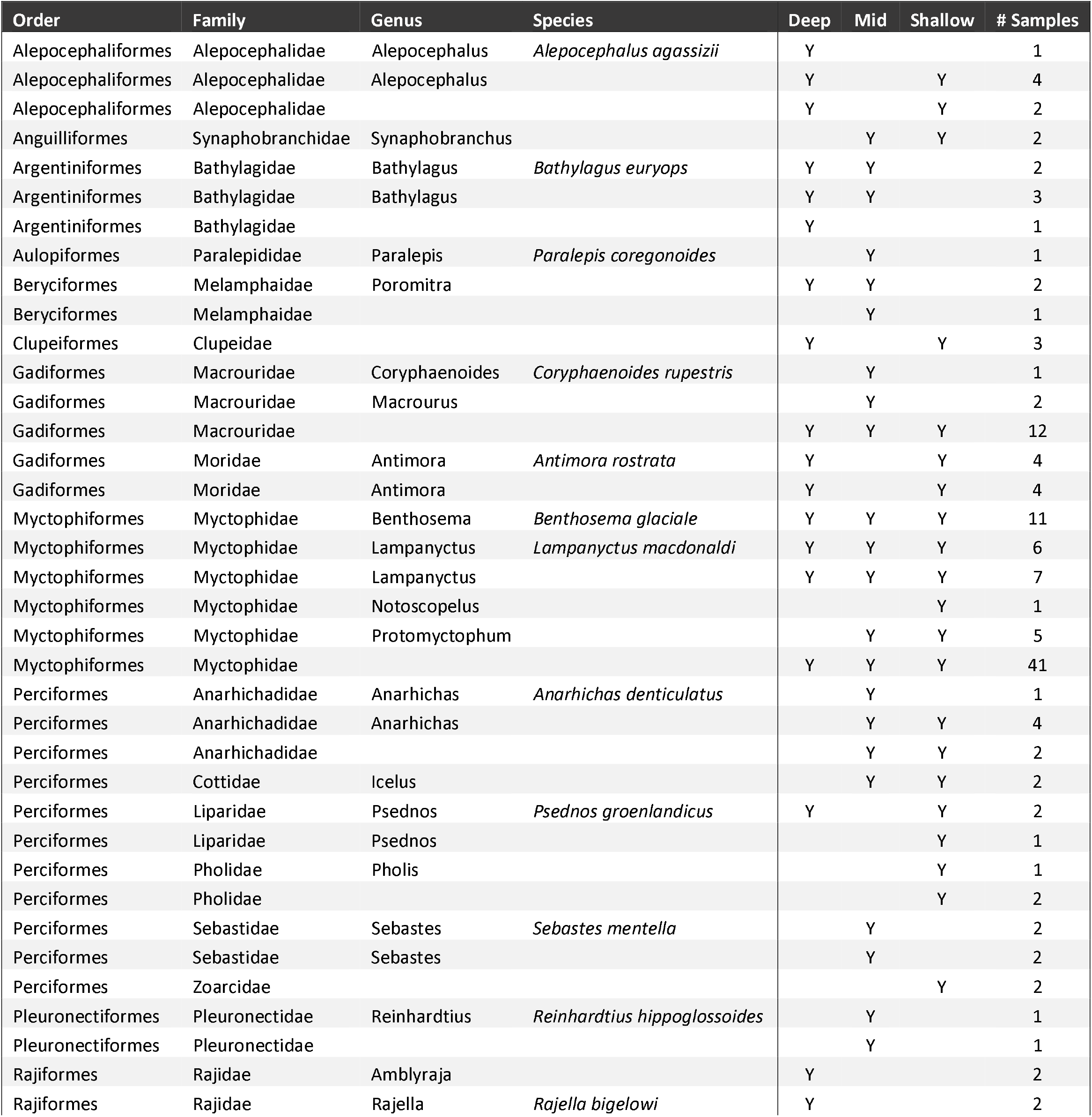
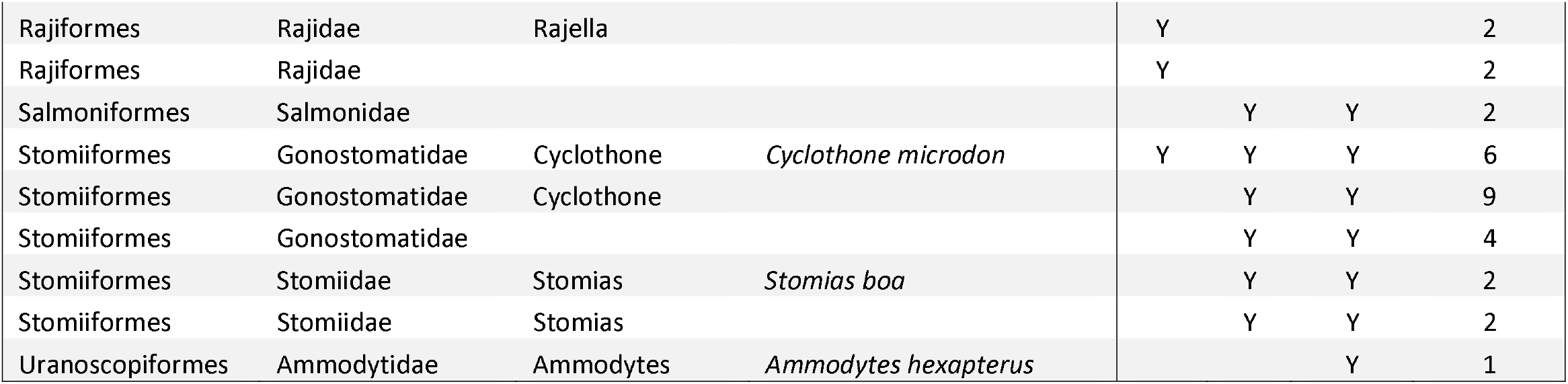
Summary of all fish taxa identified in seawater samples, indicating whether or not the taxa was detected at each depth (shallow < 500 m, mid 500 – 1400 m, deep > 1400 m) and the total number of samples in which the taxa was detected.

A total of 21 fish families, 23 genera and 15 species were identified using eDNA from 2018 and 2019 samples across all markers (Table 3). In the deep-water samples (>1400 m), 11 fish families, 11 genera and 8 species were identified. The fish species detected included several deep-water and demersal specialists, such as Bigelow’s Ray (*Rajella bigelowi*), Agassiz’ Slickhead (*Alepocephalus agassizii*), Greenland Dwarf Snailfish (*Psednos groenlandicus*), along with the Roundnose Grenadier (*Coryphaenoides rupestris*) and the Northern Wolffish (*Anarhichas denticulatus*), which are listed as Critically Endangered and Endangered, respectively on the IUCN Red List [39,40]. Several globally important members of the mesopelagic community were also detected, including Glacier Lanternfish (*Benthosema glaciale*) and Veiled Anglemouth (*Cyclothone microdon*). In addition to the fishes, 13 metazoan phyla were detected, where 58 families, 39 genera, 25 species were assigned names (S2 Table).

### Volume Comparison

Based on a two-way ANOVA, there was a significant increase in the total amount of DNA recovered (as measured by fluorometry of DNA extracts) from the 1.5-liter samples collected in 2019 compared to the 250 mL samples from 2018 (F = 219.32, df = 1, *p* < 0.001; Figure 1A). Water depth also had a significant effect on the amount of DNA recovered (F = 35.64, df =2, *p* < 0.001), with a lower DNA concentration recovered from deep water samples (>1400 m) compared to mid-depth (500-1400 m) and shallow (<500 m) samples.

**Fig 1.**
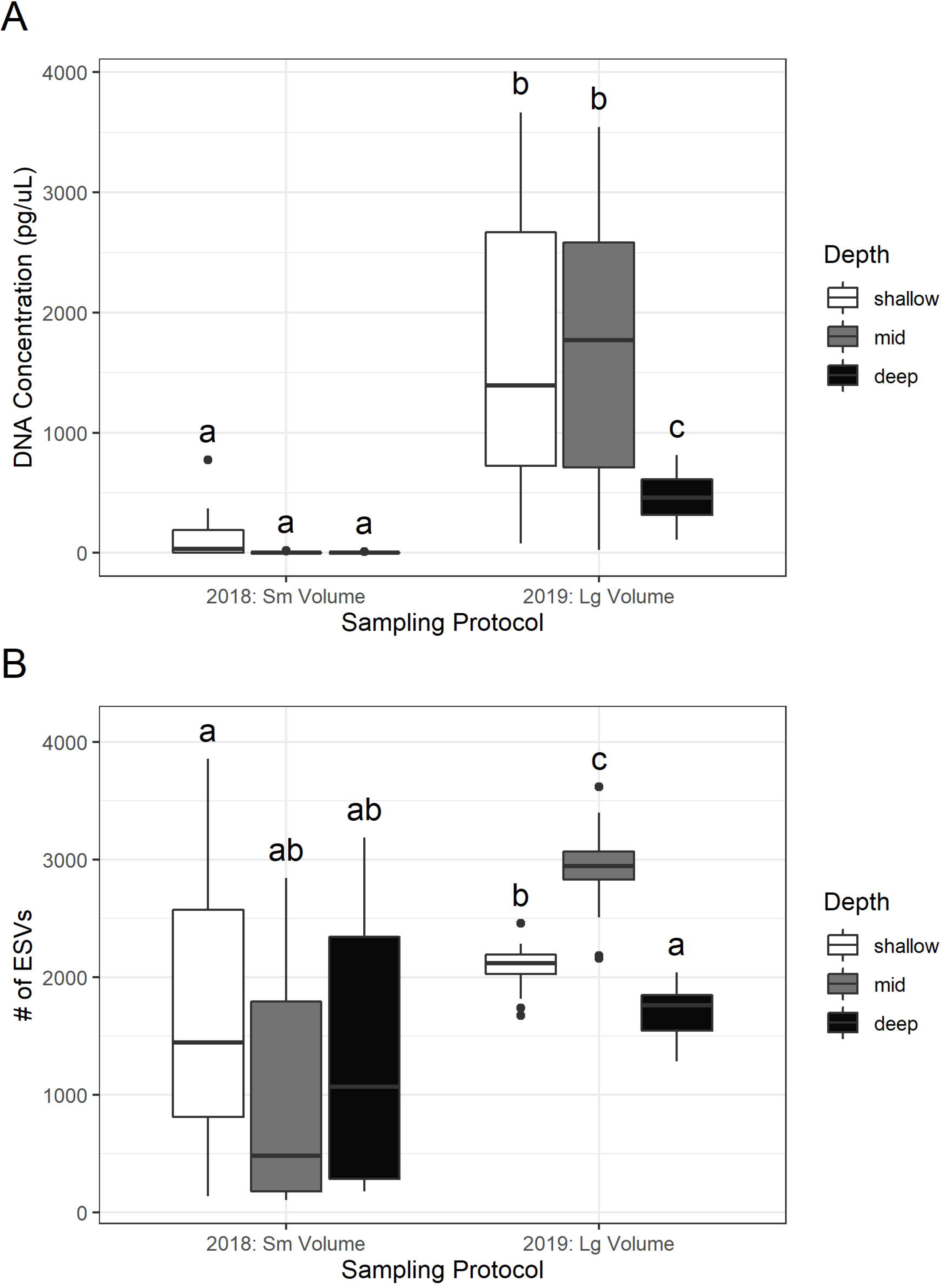
Comparison of (A) DNA concentration (pg/μL) in extracts and (B) number of ESVs recovered from small volume samples collected in 2018 and large volume samples collected in 2019 at various depths (shallow < 500 m, mid 500-1400 m, deep >1400 m). The lines inside the boxes represents the median values, the top and bottom of the boxes represent the 75% and 25% quartiles. The whiskers represent 1.5 times the inter-quartile range (IQR). Outliers (any data beyond 1.5*IQR) are shown by circles. Different letters indicate significant differences.

Based on a two-way ANOVA, there were significantly more ESVs detected in the 1.5-liter 2019 samples compared to the 250 mL 2018 samples (F = 88.28, df = 1, *p* < 0.001; Figure 1B). Additionally, there was significantly less variance in the number of ESVs recovered from the larger volume samples (F = 30.00, df = 1, *p* < 0.001). The different sampling volumes were collected in different years and at different locations so no direct comparisons of the biodiversity detected by volume could be made. There was a significant difference in the number of ESVs recovered between sampling depths (F = 6.53, df = 2, *p* = 0.002). Post-hoc comparisons revealed that large volume mid depth samples from 2019 recovered the most ESVs. The number of ESVs recovered from large volume deep samples in 2019 was not significantly higher than the number of ESVs detected in any of the small volume water samples. See S3 Fig for a comparison between DNA concentration and number of ESVs by sampling location (surface, deep scattering layer, bottom).

### Marker Comparison

Of the seven primers sets tested on the 2018 samples, the fish-targeted 12Steleo, 12S MiFishU and COI MiniFishE primer sets were the most effective at detecting fish with 11, 8 and 3 families detected by each primer set respectively. 12Steleo also provided the highest resolution with 7 species identified. COI Leray and COI F230 failed to detect any fish families. The two primer sets that identified the most metazoan families other than fish, were 18SV9M and COI FishE, which detected 18 and 12 families in 2018 samples, respectively. The three primer sets which performed well for fishes were run on samples from 2019 (12Steleo, 12S MiFishU and COI FishE) to maximize fish detection while also identifying a range of metazoans. An additional three fish families and 22 metazoan families were detected in the 2019 samples.

For the three primer sets used across both years, no single fish species was detected by all primer sets and all three primer sets detected at least one species that was unique to that primer set. Occupancy modeling revealed taxa specific variability in probabilities of detection between primer sets at the species and family level (Figure 2 & 3), however when comparing the primer sets across the whole fish community, there was little difference in the community mean probability of detection for each primer set (Figure 4).

**Fig 2.**
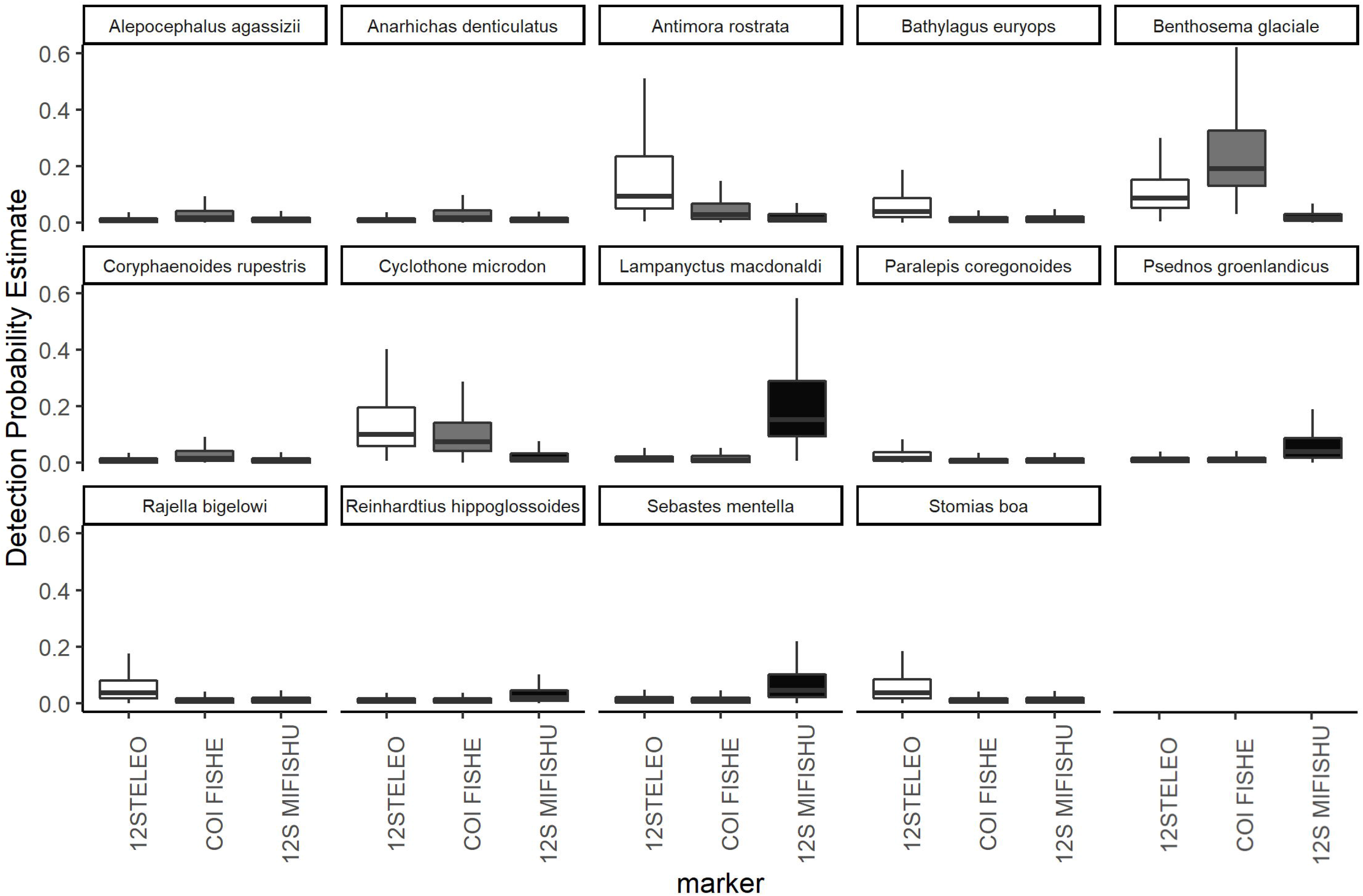
Estimated detection probability for each fish species with each primer set based on multi-species, multi-scale occupancy modeling. The lines inside the boxes represents the median values, the top and bottom of the boxes represent the 75% and 25% quartiles. The whiskers represent 1.5 times the inter-quartile range (IQR).

**Fig 3.**
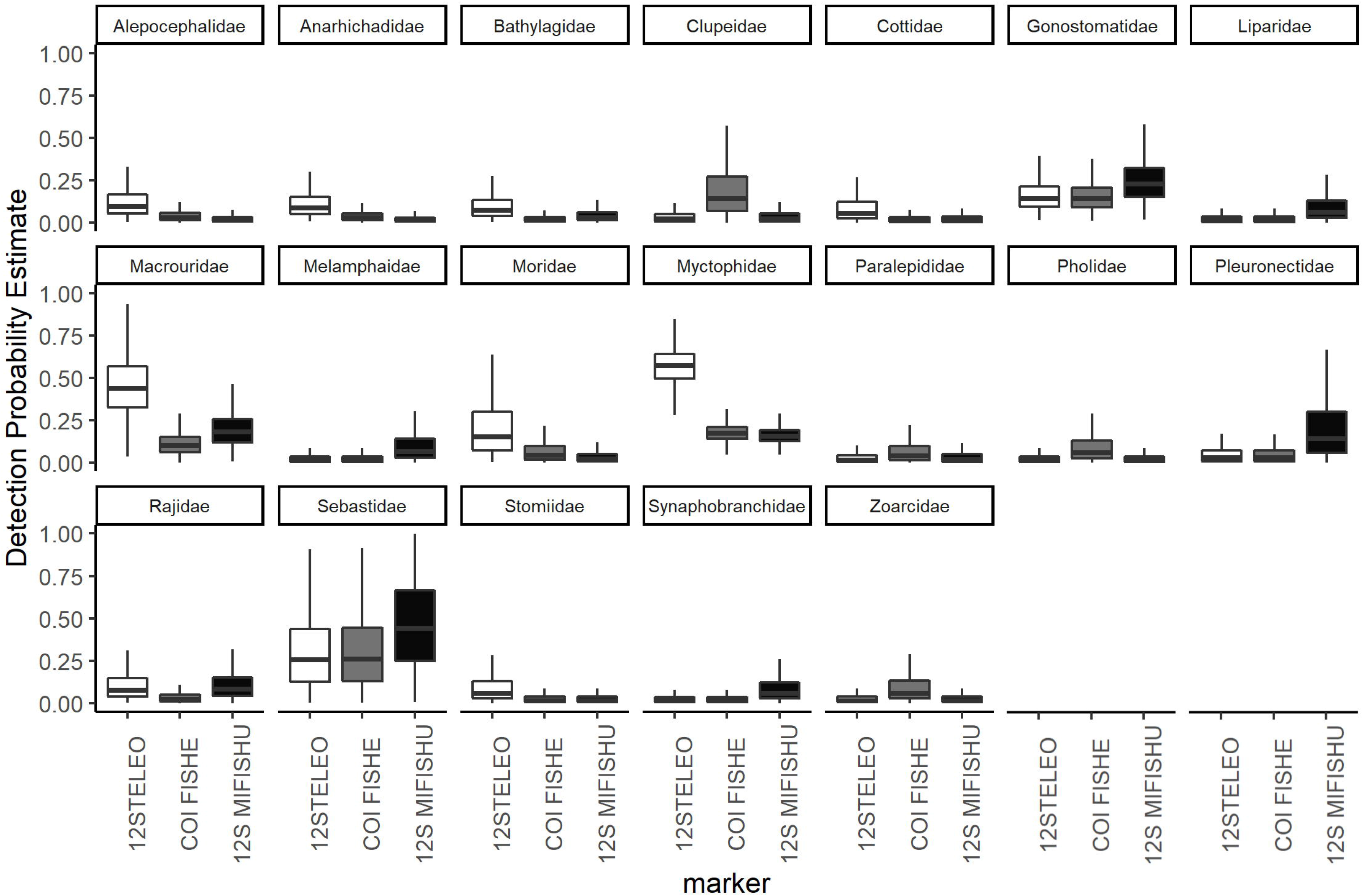
Estimated detection probability for each fish family with each primer set based on multi-species, multi-scale occupancy modeling. The lines inside the boxes represents the median values, the top and bottom of the boxes represent the 75% and 25% quartiles. The whiskers represent 1.5 times the inter-quartile range (IQR).

**Fig 4.**
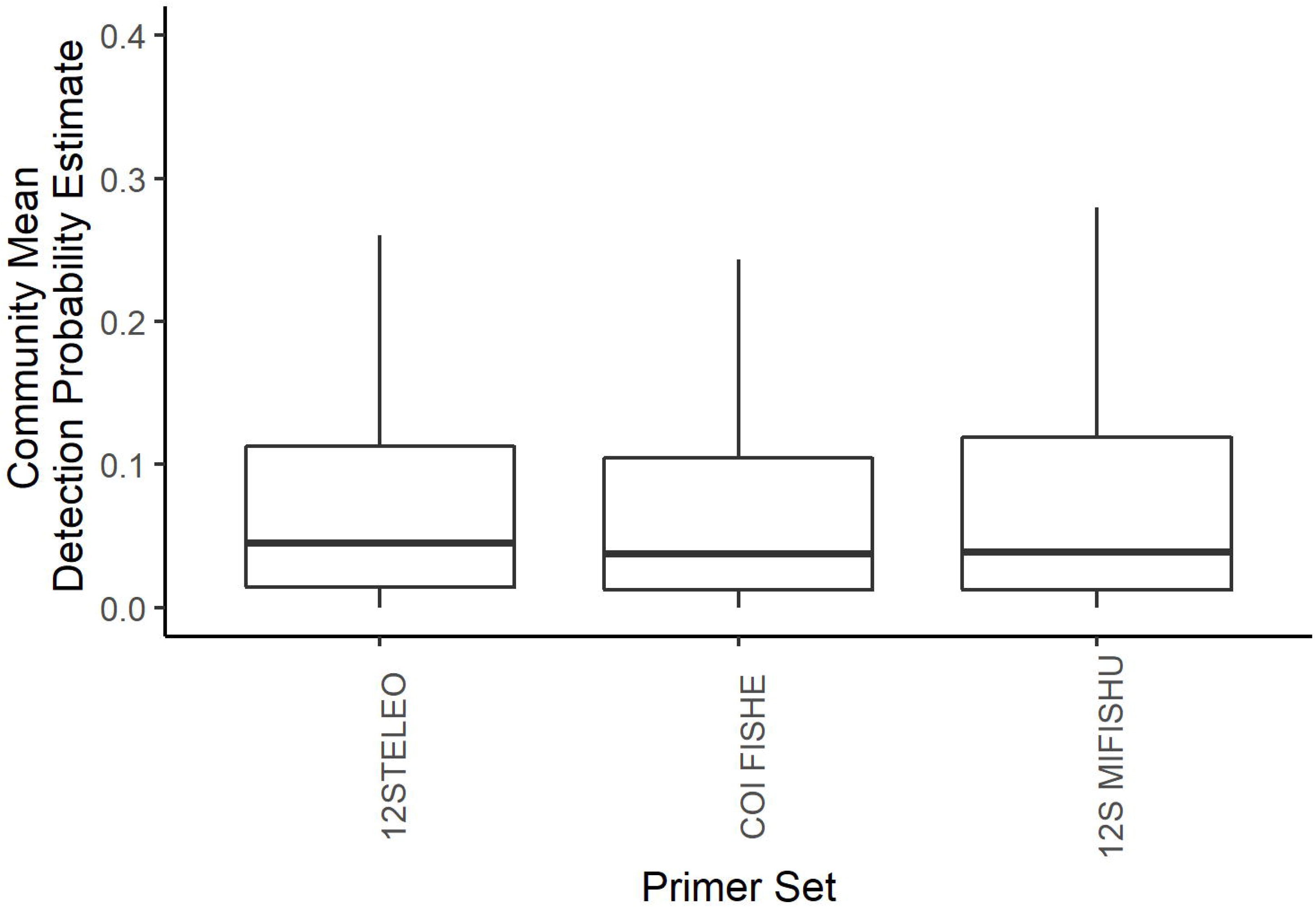
Community mean probabilities of detection for each primer set based on a multi-species, multi-scale occupancy model using fish family level data only. Similar results were seen from the fish species-level model. The lines inside the boxes represents the median values, the top and bottom of the boxes represent the 75% and 25% quartiles. The whiskers represent 1.5 times the inter-quartile range (IQR).

### Morphology & eDNA Comparison

Conventional surveys were conducted using multiple methods on three transects in the sampling area. These surveys were performed over multiple years and on multiple sampling expeditions and allowed us to assemble an inventory of species for the region. Overall, these morphological surveys identified 27 species, 25 genera and 18 families in the sampling area. To directly compare between conventional methods and eDNA, we considered only morphological data that was generated during the same sampling expedition as eDNA sample collections. There was a high degree of overlap in taxa detected between eDNA and identified via morphology (i.e. via IKMT pelagic trawls), but several taxa were unique to metabarcoding (Figure 5). A total of 14 fish species, 21 genera and 16 families were identified using eDNA while 10 fish species, 8 genera and 6 families were identified morphologically.

**Fig 5.**
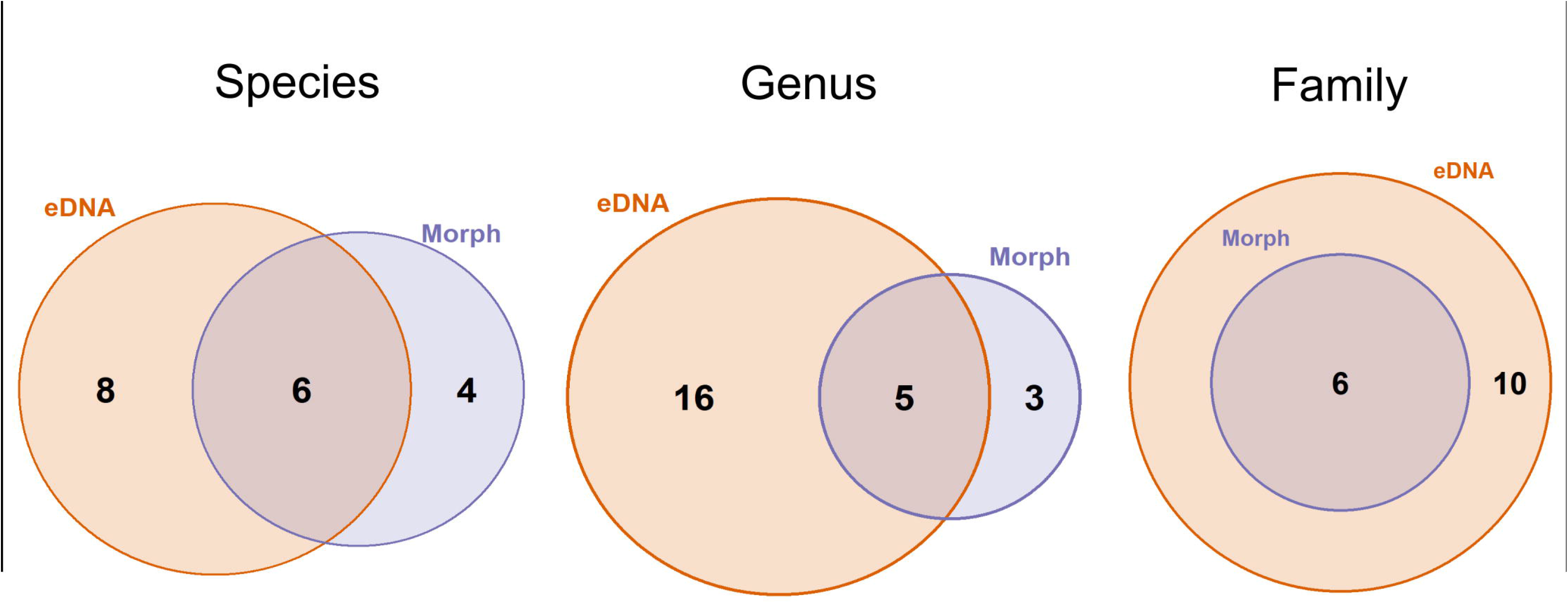
Comparison of the number of fish taxa detected at various taxonomic levels (species, genus, family) between sampling methods (eDNA metabarcoding vs. capture and morphological identification using IKMT pelagic trawls) for a single sampling expedition in 2019. Conventional methods are shown in purple and eDNA is shown in orange.

## Discussion

We demonstrated a successful protocol for the detection of deep-sea fishes using eDNA from seawater samples collected at depths down to 2500 m. Our results suggest that eDNA is less abundant in seawater from depths > 1400 m, a factor which should be considered for sampling designs of future deep-sea eDNA studies. The physical characteristics of the deep ocean (e.g. lower temperature, less sunlight) suggest DNA persists longer in this environment than at the surface [41,42], however the lower DNA concentration may reflect the different biological community present, with less abundant plankton and more species with low metabolic rates living in the deep ocean [18,43,44]. We recommend the sampling protocol followed in 2019 where larger water volumes (≥ 1.5 L) were collected, particularly for sampling the deep marine environment where the amount of DNA recovered from samples was lower. While the number of ESVs recovered from large volume deep samples was not significantly higher than the small volume samples, the reduced variance in the number of ESVs recovered suggests a more robust sampling method. Increasing the sequencing depth may be a means to make up for low DNA recovery in samples such as this. Indeed, in this study, the samples were sequenced at much higher depth (~1,000,000 reads per sample per amplicon) than most metabarcoding studies [22]. Despite an equally high sequencing depth in the small volume and low DNA concentration samples, the number of ESVs recovered was consistently higher in large volume and high DNA concentration samples. These results highlight the need for metabarcoding sampling methods to be tailored to the sampling environment and for further research into the origin, persistence, and degradation of eDNA in marine systems. Much of the research on the dynamics of eDNA has focused on freshwater systems (e.g. [45–47]) and much less is known about this cycle for eDNA in marine environments, particularly in the deep ocean (but see [48,49]). As our understanding of eDNA dynamics in the deep ocean progresses, eDNA can be a reliable way of detecting deep sea organisms, provided appropriate sampling methods are used.

When field sampling protocols are optimized for a particular system, there are often logistical constraints that must be considered in addition to the biological factors. In this study, the cold storage of large volume water samples on the sampling vessel was a limitation and therefore large volume samples were filtered *in situ* on the vessel rather than in a dedicated pre-PCR lab where downstream processing occurred. While this allowed larger water volumes to be collected, it required additional personnel time on the vessel and there may have been an increased risk of contamination for filtering *in situ* on an operational vessel at sea. In this case, precautions were taken to minimize the contamination risk including decontaminating the lab and the addition of negative controls at every step in the field (sample collection, filtration) and subsequent laboratory steps (extraction, PCR amplification). The adaptability of the filtering process was essential for allowing the collection of large volume water samples in this study. We acknowledge that these additional changes to the protocol may have contributed to the different results seen in 2018 and 2019, however water sampling volume is known to affect the biodiversity recovered from metabarcoding samples [50] and was likely the primary factor contributing to the observed differences between study years. The specific logistical constraints of sampling will be unique to each sampling mission and depend on the resources available, but they are an important consideration when optimizing sampling protocols.

We identified multiple primers sets that performed well for deep-sea fishes, but we also determined that these primer sets vary considerably in their detection probabilities within the fishes. It should also be noted that the fish-specific primers used in this study (12Steleo, 12S MiFishU) were designed to target bony fish and not cartilaginous fish. While we did detect one species of cartilaginous fish (*Rajella bigelowi*) using 12Steleo, alternative primers should be considered for studies targeting cartilaginous fish (e.g. 12S MiFishE [29]). The fish primer sets used in this study recovered many fish taxa, however the species-level resolution was not always consistently high. For example, for the 12Steleo primer set, 110 fish ESVs were recovered and only 7 (6.4%) were identified to the species level (as seen in Table 2). The low resolution is due to a combination of low sequence diversity between species (where query sequences matched multiple species in the reference database) and poor reference database coverage (where query sequences did not match any reference sequences at our species-level threshold) [51]. This reinforces the importance of marker selection and highlights the need to use multiple markers to maximize detection and taxonomic resolution even within a relatively narrow target group, such as fish. Integrating data from multiple primer sets from multiple marker regions is often recommended for metabarcoding-based biodiversity surveys [52–55]. This also highlights the need for improved species coverage in reference databases. We also identified a primer set (COI MiniFishE [24]) that performs well for a range of metazoan taxa in addition to fishes, suggesting this would be a useful primer set for comprehensive biodiversity assessments in marine environments. Conversely, one of the primer sets that has been used in a number of marine metabarcoding papers (COI Leray; mlCOIintF/ jgHCO2198 [23]) did not detect any fish taxa and hence is not recommended for analyses of fish biodiversity. Using deep sequencing with multiple primer sets is a simple strategy that can capture deep sea biodiversity especially for less abundant and elusive fish taxa.

The fish taxa detected using eDNA metabarcoding were comparable to those identified via conventional fish survey methods, although several taxa were unique to each method. This is consistent with other studies comparing eDNA to other methods of biodiversity assessment (e.g. [56,57]). When looking at a single sampling expedition, eDNA captured more fish diversity than conventional methods, and did so from rosette deployments that were used to fulfil other mission objectives (i.e. water sampling). Given the expense and time constraints associated with large research vessels, achieving such efficiencies is noteworthy. Furthermore, the relative simplicity of eDNA sample collection allows for synchronous usage of hydroacoustics and *in-situ* sensors. Metabarcoding also has the added benefit of potentially detecting species outside the target taxa. While this is dependent on the primer sets selected, the ability to detect species from all trophic levels and life histories from the same sample drastically increases the efficiency of biodiversity assessments by minimizing the number of different sampling methods required to holistically survey an ecosystem (e.g. [58]). Furthermore, various marine habitats (e.g. pelagic, demersal) can be sampled using the same methods compared to conventional surveys where different capture methods and their associated biases are used in each habitat. And finally, eDNA samples, once collected, can be used for subsequent analyses with other primer sets to generate biodiversity data for other groups or to target specific species or their populations without the need for additional sampling campaigns. For example, while the samples used in this study were collected and processed with the goal of detecting fishes, these same water samples could be processed with primers targeting corals to provide insight into deep-sea coral diversity without the need for additional sampling effort.

While there is a lot to be gained by applying metabarcoding tools to surveying the deep ocean, there are also limitations to this method. Since the biodiversity of this environment is not well-known, the reference database coverage for deep sea species is unlikely to be as comprehensive as coastal or freshwater systems. Low reference database coverage can reduce the taxonomic resolution of eDNA studies [59]. This limitation can be dealt with by generating a reference library for key fish species in the deep ocean alongside eDNA metabarcoding monitoring efforts. Metabarcoding is also limited in its quantitative ability [60] and most studies use a presence/absence approach (e.g. [61]). This method is very useful for assessing species richness and community structure [62], and determining species distributions [63], but the current methodology cannot be used to infer absolute abundance. Age structure, reproductive stage, and contaminant load are other examples of data that cannot be determined via eDNA. These factors will still rely on the capture of specimens, however eDNA can significantly increase our understanding of spatial and temporal distribution of species, which can be used to guide more detailed sampling where conventional sampling is required.

eDNA metabarcoding is a powerful approach for surveying biodiversity in the deep ocean. While future work will continue to improve these methods, such as increasing the taxonomic coverage in reference databases and refining sampling designs, this methodology can be employed immediately to complement ongoing biodiversity monitoring efforts in the deep ocean. Given the vastness of the deep ocean environment, our limited knowledge of this region’s biodiversity and the increasing anthropogenic pressures facing this fauna, there is huge potential for eDNA metabarcoding to revolutionize biodiversity monitoring and environmental stewardship in these areas.

## Acknowledgements

We would like to thank Kerry Hobrecker for her assistance in sample processing at CEGA and Jasmin Schuster, Andrew Murphy, Amy McAllister, Thibaud DeZutter, Sheena Roul, Jennica Seiden and Catie Young for their assistance processing samples at sea. Samples used for this study were collected from the Canadian research icebreaker CCGS Amundsen with the support of the Amundsen Science program funded by the Canada Foundation for Innovation (CFI) Major Science Initiatives (MSI) Fund.

## Supporting Information

**S1 Fig. Map of the sampling area in the Labrador Sea showing sampling sites along three transects that follow a depth gradient of approximately 500 m to 3000 m.** Colours of sampling sites indicate the year of eDNA sampling. Inset map shows the location of the sampling area on a global map. Map data source: Esri. Ocean Reference [basemap]. 1:6000000. Ocean Basemap. February 10, 2012. www.arcgis.com/home/item.html?id=5ae9e138a17842688b0b79283a4353f6. (Accessed: June 15, 2020).

**S2 Fig. Scatterplot plot comparing water sampling depth and DNA concentration for eDNA water samples collected in the Labrador Sea in 2019.** The blue line represents the predicted values based on a generalized linear model with 95% confidence intervals shown in gray.

**S3 Fig. Comparison of (A) DNA concentration (pg/μL) in extracts and (B) number of ESVs recovered from small volume samples collected in 2018 and large volume samples collected in 2019 at various depth sampling locations (surface, deep scattering layer, bottom).** Different letters indicate significant differences.

**S1 Text. Detailed occupancy modeling methods and model structure.**

**S1 Table. Sampling summary table listing the sampling stations, their location, date of collection, sampling depths and approximate water depth.** Triplicate water samples were collected at each station and date listed.

**S2 Table. Summary of all metazoan taxa identified in seawater samples, indicating whether or not the taxa was detected at each depth (shallow < 500 m, mid 500 – 1400 m, deep > 1400 m) and the total number of samples in which the taxa was detected.**

