## Supplementary material for "Harnessing the power of eDNA metabarcoding for the detection of deep-sea fishes": S1 Fig

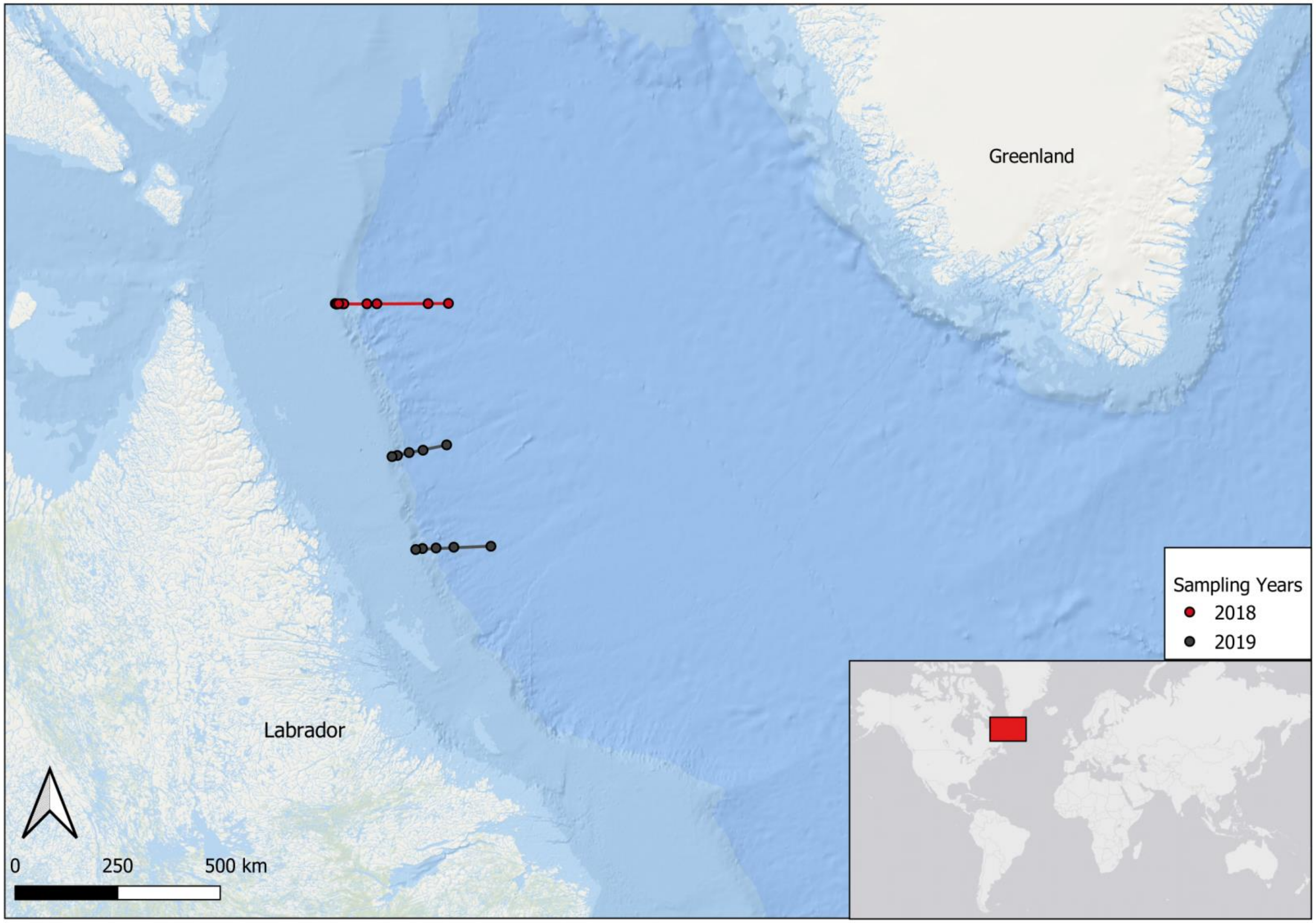


**S1 Fig. Map of sampling area in the Labrador Sea showing sampling sites along three transects that follow a depth gradient of approximately 500 m to 3000 m.** Colours of sampling sites indicate the year of eDNA sampling. Inset map shows the location of the sampling area on a global map. Map data source: Esri. Ocean Reference [basemap]. 1:6000000. Ocean Basemap. February 10, 2012. www.arcgis.com/home/item.html?id=5ae9e138a17842688b0b79283a4353f6. (Accessed: June 15, 2020).
