## Supplementary material for "Harnessing the power of eDNA metabarcoding for the detection of deep-sea fishes": S1 Table

**S1 Table.** **Sampling summary table listing the sampling stations, their location, date of collection, sampling depths and approximate water depth.** Triplicate water samples were collected at each station and date listed.

| **Station** | **Year** | **Collection Date** | **Water Depth** | **Sample Depth** | **Depth Category** | **Latitude** | **Longitude** |
| --- | --- | --- | --- | --- | --- | --- | --- |
| ISECOLD_2_2500 | 2019 | 6/29/2019 | 2500 | 0 | shallow | 58.9069 | -58.8514 |
| ISECOLD_2_2500 | 2019 | 6/29/2019 | 2500 | 500 | mid | 58.9069 | -58.8514 |
| ISECOLD_2_2500 | 2019 | 6/29/2019 | 2500 | 2395 | deep | 58.9069 | -58.8514 |
| ISECOLD_2_2000 | 2019 | 6/29/2019 | 2000 | 0 | shallow | 58.8468 | -59.3684 |
| ISECOLD_2_2000 | 2019 | 6/29/2019 | 2000 | 500 | mid | 58.8468 | -59.3684 |
| ISECOLD_2_2000 | 2019 | 6/29/2019 | 2000 | 1937 | deep | 58.8468 | -59.3684 |
| ISECOLD_2_1000 | 2019 | 6/30/2019 | 1000 | 0 | shallow | 58.7869 | -59.9300 |
| ISECOLD_2_1000 | 2019 | 6/30/2019 | 1000 | 500 | mid | 58.7869 | -59.9300 |
| ISECOLD_2_1000 | 2019 | 6/30/2019 | 1000 | 1038 | mid | 58.7869 | -59.9300 |
| ISECOLD_1_2500 | 2019 | 6/28/2019 | 2500 | 0 | shallow | 57.7406 | -57.8814 |
| ISECOLD_1_2500 | 2019 | 6/28/2019 | 2500 | 500 | mid | 57.7406 | -57.8814 |
| ISECOLD_1_2500 | 2019 | 6/28/2019 | 2500 | 2492 | deep | 57.7406 | -57.8814 |
| ISECOLD_1_2000 | 2019 | 6/27/2019 | 2000 | 1981 | deep | 57.7296 | -58.6936 |
| ISECOLD_1_1500 | 2019 | 6/26/2019 | 1500 | 0 | shallow | 57.7202 | -59.0831 |
| ISECOLD_1_1500 | 2019 | 6/26/2019 | 1500 | 500 | mid | 57.7202 | -59.0831 |
| ISECOLD_1_1500 | 2019 | 6/26/2019 | 1500 | 1474 | deep | 57.7202 | -59.0831 |
| ISECOLD_1_1000 | 2019 | 6/25/2019 | 1000 | 1010 | mid | 57.7140 | -59.3796 |
| ISECOLD_1_500 | 2019 | 6/25/2019 | 500 | 0 | shallow | 57.7022 | -59.5283 |
| ISECOLD_1_500 | 2019 | 6/25/2019 | 500 | 600 | mid | 57.7022 | -59.5283 |
| ISECOLD_2_500 | 2019 | 6/30/2019 | 500 | 0 | shallow | 58.7743 | -60.0475 |
| ISECOLD_2_500 | 2019 | 6/30/2019 | 500 | 527 | mid | 58.7743 | -60.0475 |
| ISECOLD_2_1500 | 2019 | 6/30/2019 | 1500 | 0 | shallow | 58.8196 | -59.6732 |
| ISECOLD_2_1500 | 2019 | 6/30/2019 | 1500 | 500 | mid | 58.8196 | -59.6732 |
| ISECOLD_2_1500 | 2019 | 6/30/2019 | 1500 | 1496 | deep | 58.8196 | -59.6732 |
| DFO-1 | 2019 | 7/2/2019 | 500 | 500 | mid | 60.4691 | -61.2903 |
| DFO-3 | 2019 | 7/2/2019 | 1000 | 1000 | mid | 60.4678 | -61.1460 |
| DFO-1 | 2018 | 7/29/2018 | 500 | 506 | mid | 60.4635 | -61.2645 |
| DFO-1 | 2018 | 7/29/2018 | 500 | 396 | shallow | 60.4635 | -61.2645 |
| DFO-1 | 2018 | 7/29/2018 | 500 | 3 | shallow | 60.4635 | -61.2645 |
| DFO-3 | 2018 | 7/31/2018 | 1000 | 1122 | mid | 60.4663 | -61.1041 |
| DFO-3 | 2018 | 7/31/2018 | 1000 | 594 | mid | 60.4663 | -61.1041 |
| DFO-3 | 2018 | 7/31/2018 | 1000 | 3 | shallow | 60.4663 | -61.1041 |
| DFO-3 | 2018 | 7/31/2018 | 1000 | 250 | shallow | 60.4663 | -61.1041 |
| DFO-750 | 2018 | 7/31/2018 | 750 | 706 | mid | 60.4672 | -61.2177 |
| DFO-750 | 2018 | 8/1/2018 | 750 | 497 | shallow | 60.4672 | -61.2177 |
| DFO-750 | 2018 | 8/1/2018 | 750 | 2 | shallow | 60.4672 | -61.2177 |
| DFO-750 | 2018 | 8/1/2018 | 750 | 247 | shallow | 60.4672 | -61.2177 |
| DFO-5 | 2018 | 8/2/2018 | 1500 | 1418 | deep | 60.4669 | -60.5977 |
| DFO-5 | 2018 | 8/2/2018 | 1500 | 501 | mid | 60.4669 | -60.5977 |
| DFO-5 | 2018 | 8/2/2018 | 1500 | 2 | shallow | 60.4669 | -60.5977 |
| DFO-5 | 2018 | 8/2/2018 | 1500 | 299 | shallow | 60.4669 | -60.5977 |
| DFO-7 | 2018 | 8/2/2018 | 2000 | 1878 | deep | 60.4669 | -60.3800 |
| DFO-7 | 2018 | 8/2/2018 | 2000 | 495 | shallow | 60.4669 | -60.3800 |
| DFO-7 | 2018 | 8/2/2018 | 2000 | 3 | shallow | 60.4669 | -60.3800 |
| DFO-8 | 2018 | 8/3/2018 | 2500 | 2428 | deep | 60.4685 | -59.2575 |
| DFO-9 | 2018 | 8/3/2018 | 2500 | 2502 | deep | 60.4710 | -58.8132 |
| DFO-9 | 2018 | 8/4/2018 | 2500 | 496 | shallow | 60.4710 | -58.8132 |
| DFO-9 | 2018 | 8/4/2018 | 2500 | 2 | shallow | 60.4710 | -58.8132 |
