## Supplementary material for "Harnessing the power of eDNA metabarcoding for the detection of deep-sea fishes": S1 Text

**S1 Text. Detailed occupancy modeling methods and model structure.**

**Occupancy Modeling**

Under the occupancy modeling framework described in ^1^, each sampling depth at each station was considered a different site in the occupancy model. Replicate bottles collected at a site were considered samples. Each of the three primer sets (used in 2018 and 2019; 12Steleo, MiFishU, FishE) amplified from each bottle was considered a technical replicate.

We included primer set as a categorical covariate (α1) at the level of probability of detection. We included water depth (m) as a continuous covariate (α2) at the level of occupancy. Covariates were included in the model as follows:

$$logit\left( p_{ijrk} \right)= {lp}_{k}+ {\beta1}_{k {\alpha1}_{ijr}}$$

$$logit\left( \psi_{ik} \right)={lpsi}_{k}+ {\beta2}_{k}* {\alpha2}_{i}$$

The water depth covariate was z-score standardized to have a mean of zero and a standard deviation of one to help with model convergence. Species coefficients arise from additional community-level parameters:

$${lpsi}_{k} \sim N(\mu_{lpsi}, \sigma_{lpsi})$$

$${ltheta}_{k} \sim N(\mu_{ltheta}, \sigma_{ltheta})$$

$${lp}_{k} \sim N(\mu_{lp}, \sigma_{lp})$$

$${\beta1}_{k} \sim N(\mu_{\beta1}, \sigma_{\beta1})$$

$${\beta2}_{k \alpha2} \sim N(\mu_{\beta2}, \sigma_{\beta2})$$

$${\beta3}_{k} \sim N(\mu_{\beta3}, \sigma_{\beta3})$$

Community-level parameters were described by weakly informative hyperpriors ^2^. All mean values for the above prior distributions were selected from a normal distribution and all standard deviations were selected from a uniform distribution.

$$\mu\sim N(0,10)$$

$$\sigma\sim Uniform(0,5)$$

All statistical analyses were conducted in R v3.5.1 ^3^. MCMC sampling was achieved with JAGS ^4^, implemented using ‘jagsUI’ v1.5.0 ^5^. The model was written for JAGS in the BUGS language (see below for model structure). For each model (at the family level and at the species level), MCMC sampling was run in three chains, each with 50,000 iterations, a burn in of 10,000, and a thinning rate of 10. Convergence was verified using the Gelman-Rubin diagnostic ^6^ and by evaluating trace plots. For both models, parameter estimates are presented in boxplot format.

**Model Structure**

model {

\# Priors for species-specific effects in occupancy and detection

for(k in 1:nspec){

lpsi[k] ~ dnorm(mu.lpsi, tau.lpsi)	# Hyperparams describe community

ltheta[k] ~ dnorm(mu.ltheta, tau.ltheta)

betalpsi2[k] ~ dnorm(mu.betalpsi2, tau.betalpsi2)

lp[k] ~ dnorm(mu.lp, tau.lp)

for (m in 1:ntechrep) {

betalp1[k,m] ~ dnorm(mu.betalp1, tau.betalp1)

}

}

\# Hyperpriors

\# For the model of occupancy

mu.lpsi ~ dnorm(0,0.01)

sd.lpsi ~ dunif(0,5) # as always, bounds of uniform chosen by trial and error

tau.lpsi <- pow(sd.lpsi, -2)

mu.betalpsi2 ~ dnorm(0,0.01)

sd.betalpsi2 ~ dunif(0, 5)

tau.betalpsi2 <- pow(sd.betalpsi2, -2)

\# For the model of availability

mu.ltheta ~ dnorm(0,0.01)

sd.ltheta ~ dunif(0, 5)

tau.ltheta <- pow(sd.ltheta, -2)

\# For the model of detection

mu.lp ~ dnorm(0,0.01)

sd.lp ~ dunif(0, 5)

tau.lp <- pow(sd.lp, -2)

mu.betalp1 ~ dnorm(0,0.01)

sd.betalp1 ~ dunif(0, 5)

tau.betalp1 <- pow(sd.betalp1, -2)

\# Ecological model for true occurrence (process model)

for(k in 1:nspec){

for (i in 1:nsite) {

logit(psi[i,k]) <- lpsi[k] + betalpsi2[k] * depth[i]

z[i,k] ~ dbern(psi[i,k])

}

}

\# Observation model for replicate sample bottles

for(k in 1:nspec){

for (i in 1:nsite){

for(j in 1:nbiorep){

logit(theta[i,j,k]) <- ltheta[k]

mu.theta[i,j,k] <- z[i,k] * theta[i,j,k]

w[i,j,k] ~ dbern(mu.theta[i,j,k])

}

}

}

\# Observation model for replicated primers for each sample

for(k in 1:nspec){

for (i in 1:nsite){

for(j in 1:nbiorep){

for (r in 1:ntechrep){

logit(p[i,j,r,k]) <- lp[k] + betalp1[k,marker[i,j,r]]

mu.p[i,j,r,k] <- w[i,j,k] * p[i,j,r,k]

y[i,j,r,k] ~ dbern(mu.p[i,j,r,k])

}

}

}

}

}
