## Supplementary material for "Harnessing the power of eDNA metabarcoding for the detection of deep-sea fishes": S2 Fig

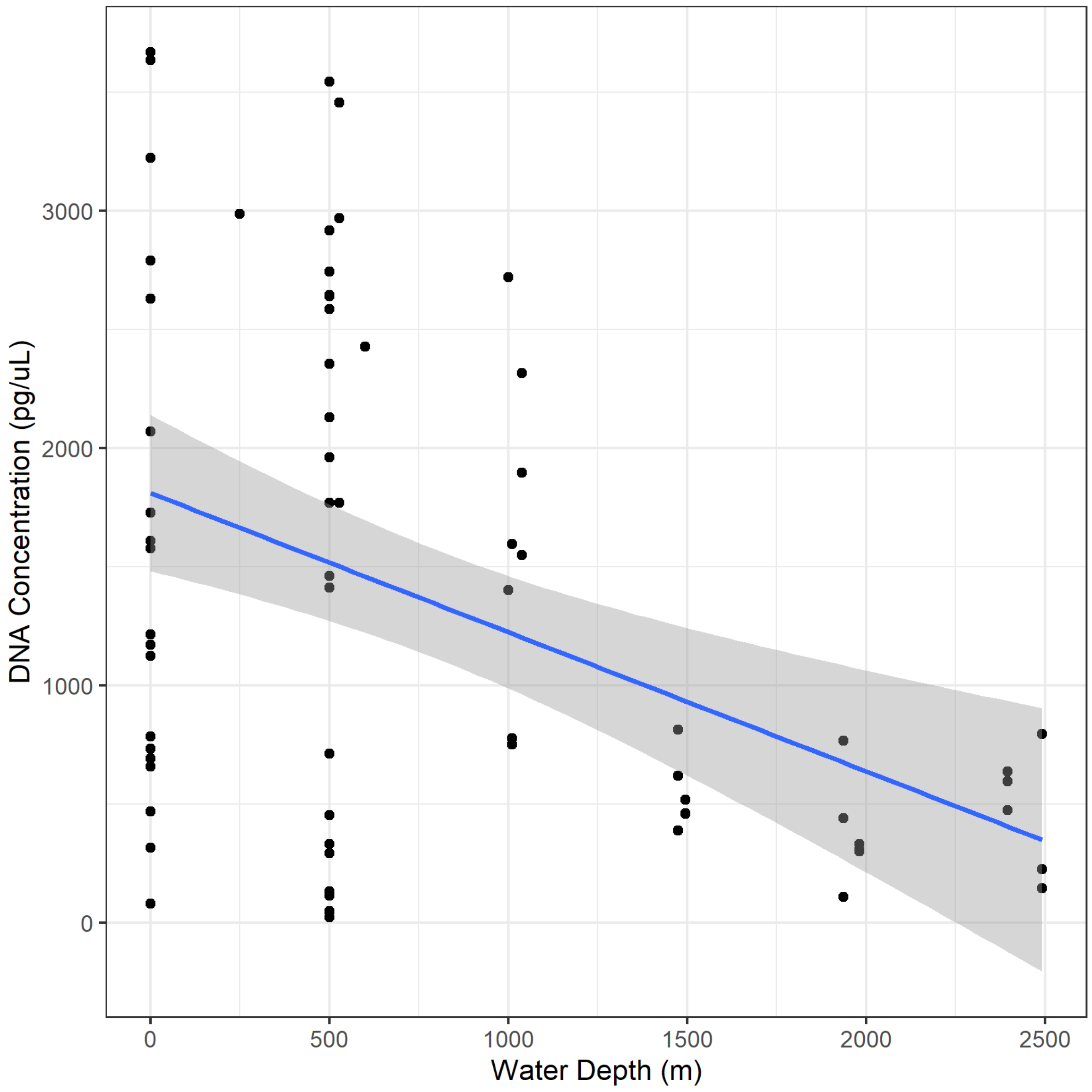


**S2 Fig. Scatterplot plot comparing water sampling depth and DNA concentration for eDNA water samples collected in the Labrador Sea in 2019.** The blue line represents the predicted values based on a generalized linear model with 95% confidence intervals shown in gray.
