## Supplementary material for "Harnessing the power of eDNA metabarcoding for the detection of deep-sea fishes": S2 Table

**S2 Table.** **Summary of all metazoan taxa identified in seawater samples, indicating whether or not the taxa was detected at each depth ( shallow < 500 m , mid 500 – 1400 m , deep > 1400 m) and the total number of samples in which the taxa was detected.**

| **Phylum** | **Class** | **Order** | **Family** | **Genus** | **Species** | **Deep** | **Mid** | **Shallow** | **# Samples** |
| --- | --- | --- | --- | --- | --- | --- | --- | --- | --- |
| Annelida | Polychaeta | Capitellida | Scalibregmatidae | Scalibregma | *Scalibregma inflatum* |  |  | Y | 1 |
| Annelida | Polychaeta | Eunicida | Onuphidae | Nothria | *Nothria conchylega* |  | Y |  | 1 |
| Annelida | Polychaeta | Flabelligerida | Flabelligeridae |  |  |  |  | Y | 2 |
| Annelida | Polychaeta | Phyllodocida | Goniadidae | Goniada |  |  | Y |  | 2 |
| Annelida | Polychaeta | Phyllodocida | Goniadidae |  |  | Y | Y | Y | 13 |
| Annelida | Polychaeta | Phyllodocida | Nephtyidae | Micronephthys | *Micronephthys minuta* |  |  | Y | 2 |
| Annelida | Polychaeta | Spionida | Cirratulidae |  |  |  | Y | Y | 3 |
| Annelida | Polychaeta | Spionida | Spionidae |  |  | Y |  |  | 1 |
| Annelida | Polychaeta | Terebellida | Ampharetidae | Ampharete |  |  |  | Y | 1 |
| Annelida | Polychaeta | Terebellida | Terebellidae |  |  | Y |  | Y | 2 |
| Annelida | Polychaeta | Terebellida | Trichobranchidae | Terebellides |  |  | Y |  | 1 |
| Annelida |  |  |  |  |  | Y | Y | Y | 23 |
| Arthropoda | Hexanauplia | Calanoida | Acartiidae | Acartia |  |  |  | Y | 3 |
| Arthropoda | Hexanauplia | Calanoida | Acartiidae |  |  | Y | Y | Y | 7 |
| Arthropoda | Hexanauplia | Calanoida | Calanidae | Calanus | *Calanus finmarchicus* | Y | Y | Y | 31 |
| Arthropoda | Hexanauplia | Calanoida | Calanidae | Calanus | *Calanus glacialis* |  | Y | Y | 14 |
| Arthropoda | Hexanauplia | Calanoida | Calanidae | Calanus |  |  |  | Y | 32 |
| Arthropoda | Hexanauplia | Calanoida | Calanidae | Neocalanus | *Neocalanus robustior* |  |  | Y | 1 |
| Arthropoda | Hexanauplia | Calanoida | Calanidae |  |  | Y | Y | Y | 35 |
| Arthropoda | Hexanauplia | Calanoida | Clausocalanidae | Pseudocalanus | *Pseudocalanus minutus* |  |  | Y | 11 |
| Arthropoda | Hexanauplia | Calanoida | Clausocalanidae | Pseudocalanus | *Pseudocalanus newmani* |  |  | Y | 2 |
| Arthropoda | Hexanauplia | Calanoida | Clausocalanidae | Pseudocalanus |  |  |  | Y | 3 |
| Arthropoda | Hexanauplia | Calanoida | Clausocalanidae |  |  |  |  | Y | 8 |
| Arthropoda | Hexanauplia | Calanoida | Euchaetidae |  |  |  | Y |  | 1 |
| Arthropoda | Hexanauplia | Calanoida | Heterorhabdidae | Heterorhabdus |  |  | Y |  | 1 |
| Arthropoda | Hexanauplia | Calanoida | Metridinidae | Metridia |  |  | Y |  | 1 |
| Arthropoda | Hexanauplia | Calanoida | Spinocalanidae | Spinocalanus | *Spinocalanus brevicaudatus* | Y | Y |  | 2 |
| Arthropoda | Hexanauplia | Calanoida | Spinocalanidae | Spinocalanus |  |  | Y |  | 1 |
| Arthropoda | Hexanauplia | Cyclopoida | Oithonidae | Oithona | *Oithona similis* |  |  | Y | 40 |
| Arthropoda | Hexanauplia | Cyclopoida | Oithonidae | Oithona |  |  |  | Y | 12 |
| Arthropoda | Hexanauplia | Cyclopoida | Oithonidae |  |  | Y | Y | Y | 41 |
| Arthropoda | Insecta | Diptera | Chironomidae |  |  |  | Y |  | 1 |
| Arthropoda | Malacostraca | Amphipoda | Hyperiidae | Themisto |  |  |  | Y | 2 |
| Arthropoda | Malacostraca | Euphausiacea | Euphausiidae | Thysanoessa | *Thysanoessa longicaudata* | Y | Y | Y | 29 |
| Arthropoda | Malacostraca | Euphausiacea | Euphausiidae | Thysanoessa |  | Y | Y | Y | 26 |
| Arthropoda | Malacostraca | Euphausiacea | Euphausiidae |  |  |  |  | Y | 1 |
| Arthropoda |  |  |  |  |  | Y | Y | Y | 42 |
| Chaetognatha | Sagittoidea | Aphragmophora | Sagittidae | Pseudosagitta | *Pseudosagitta maxima* | Y | Y |  | 3 |
| Chaetognatha | Sagittoidea | Phragmophora | Eukrohniidae |  |  |  |  | Y | 1 |
| Chaetognatha |  |  |  |  |  |  | Y | Y | 3 |
| Chordata | Ascidiacea | Stolidobranchia | Pyuridae | Microcosmus |  | Y |  |  | 1 |
| Chordata | Ascidiacea | Stolidobranchia | Pyuridae |  |  | Y |  |  | 1 |
| Chordata | Mammalia | Cetacea | Delphinidae | Globicephala | *Globicephala melas* |  | Y | Y | 8 |
| Chordata | Mammalia | Cetacea | Delphinidae | Globicephala |  |  | Y | Y | 2 |
| Chordata | Mammalia | Cetacea | Phocoenidae | Phocoena | *Phocoena phocoena* |  | Y | Y | 2 |
| Chordata | Mammalia | Cetacea | Phocoenidae |  |  |  |  | Y | 1 |
| Chordata | Mammalia | Cetacea | Ziphiidae | Hyperoodon | *Hyperoodon ampullatus* | Y |  | Y | 2 |
| Chordata | Mammalia | Cetacea | Ziphiidae | Hyperoodon |  |  |  | Y | 1 |
| Chordata |  |  |  |  |  | Y | Y | Y | 15 |
| Cnidaria | Anthozoa | Actiniaria | Actiniidae | Urticina | *Urticina eques* |  | Y |  | 1 |
| Cnidaria | Anthozoa | Actiniaria | Actinostolidae |  |  | Y | Y |  | 2 |
| Cnidaria | Anthozoa | Actiniaria | Hormathiidae | Hormathia |  |  | Y |  | 1 |
| Cnidaria | Anthozoa | Alcyonacea | Isididae | Acanella | *Acanella arbuscula* | Y |  |  | 1 |
| Cnidaria | Anthozoa | Alcyonacea | Isididae |  |  | Y |  |  | 1 |
| Cnidaria | Anthozoa | Alcyonacea | Primnoidae |  |  | Y | Y |  | 6 |
| Cnidaria | Anthozoa | Antipatharia | Schizopathidae |  |  |  | Y |  | 1 |
| Cnidaria | Anthozoa | Scleractinia | Fungiacyathidae | Fungiacyathus |  |  | Y |  | 1 |
| Cnidaria | Anthozoa | Zoantharia | Parazoanthidae | Parazoanthus |  |  | Y |  | 1 |
| Cnidaria | Hydrozoa | Narcomedusae | Aeginidae | Aeginura | *Aeginura grimaldii* | Y |  |  | 1 |
| Cnidaria | Hydrozoa | Narcomedusae | Aeginidae |  |  |  | Y |  | 1 |
| Cnidaria | Hydrozoa | Siphonophorae | Agalmatidae | Marrus | *Marrus orthocanna* |  |  | Y | 6 |
| Cnidaria | Hydrozoa | Siphonophorae | Agalmatidae |  |  |  |  | Y | 1 |
| Cnidaria | Hydrozoa | Trachymedusae | Rhopalonematidae | Pantachogon | *Pantachogon haeckeli* |  | Y |  | 1 |
| Cnidaria | Scyphozoa | Coronatae | Atollidae | Atolla | *Atolla wyvillei* |  | Y |  | 1 |
| Cnidaria | Scyphozoa | Coronatae | Periphyllidae | Periphylla | *Periphylla periphylla* |  | Y | Y | 2 |
| Cnidaria | Scyphozoa | Semaeostomeae | Cyaneidae | Cyanea |  |  |  | Y | 1 |
| Cnidaria |  |  |  |  |  |  | Y | Y | 20 |
| Ctenophora | Tentaculata | Cydippida | Lampeidae | Lampea |  |  | Y | Y | 12 |
| Ctenophora | Tentaculata | Cydippida | Lampeidae |  |  |  |  | Y | 5 |
| Echinodermata | Asteroidea | Paxillosida | Astropectinidae |  |  |  | Y | Y | 12 |
| Echinodermata | Echinoidea | Echinoida | Strongylocentrotidae | Strongylocentrotus | *Strongylocentrotus pallidus* |  |  | Y | 1 |
| Echinodermata | Echinoidea | Echinothurioida | Echinothuriidae |  |  | Y |  |  | 1 |
| Echinodermata | Holothuroidea | Elasipodida | Elpidiidae | Peniagone |  | Y |  |  | 1 |
| Echinodermata | Ophiuroidea | Ophiurida | Ophiactidae | Ophiopholis | *Ophiopholis aculeata* |  |  | Y | 2 |
| Echinodermata | Ophiuroidea | Ophiurida | Ophiactidae | Ophiopholis |  |  |  | Y | 1 |
| Echinodermata |  |  |  |  |  |  | Y |  | 1 |
| Mollusca | Gastropoda | Littorinimorpha | Lacunidae |  |  | Y |  |  | 2 |
| Mollusca | Scaphopoda | Dentaliida | Dentaliidae |  |  |  | Y |  | 1 |
| Mollusca |  |  |  |  |  | Y | Y | Y | 13 |
| Nemertea | Enopla | Monostilifera | Oerstediidae | Oerstedia | *Oerstedia dorsalis* |  |  | Y | 1 |
| Porifera | Demospongiae | Poecilosclerida | Crellidae | Crella |  | Y | Y |  | 2 |
| Porifera | Demospongiae | Poecilosclerida | Crellidae |  |  | Y |  |  | 1 |
| Porifera | Demospongiae | Tetractinellida | Tetillidae | Craniella |  |  | Y |  | 1 |
| Porifera |  | Lyssacinosida | Rossellidae |  |  | Y |  |  | 1 |
| Porifera |  |  |  |  |  | Y | Y | Y | 7 |
| Xenacoelomorpha | | Acoela | Convolutidae |  |  |  |  | Y | 2 |
