## Supplementary material for "Harnessing the power of eDNA metabarcoding for the detection of deep-sea fishes": S3 Fig

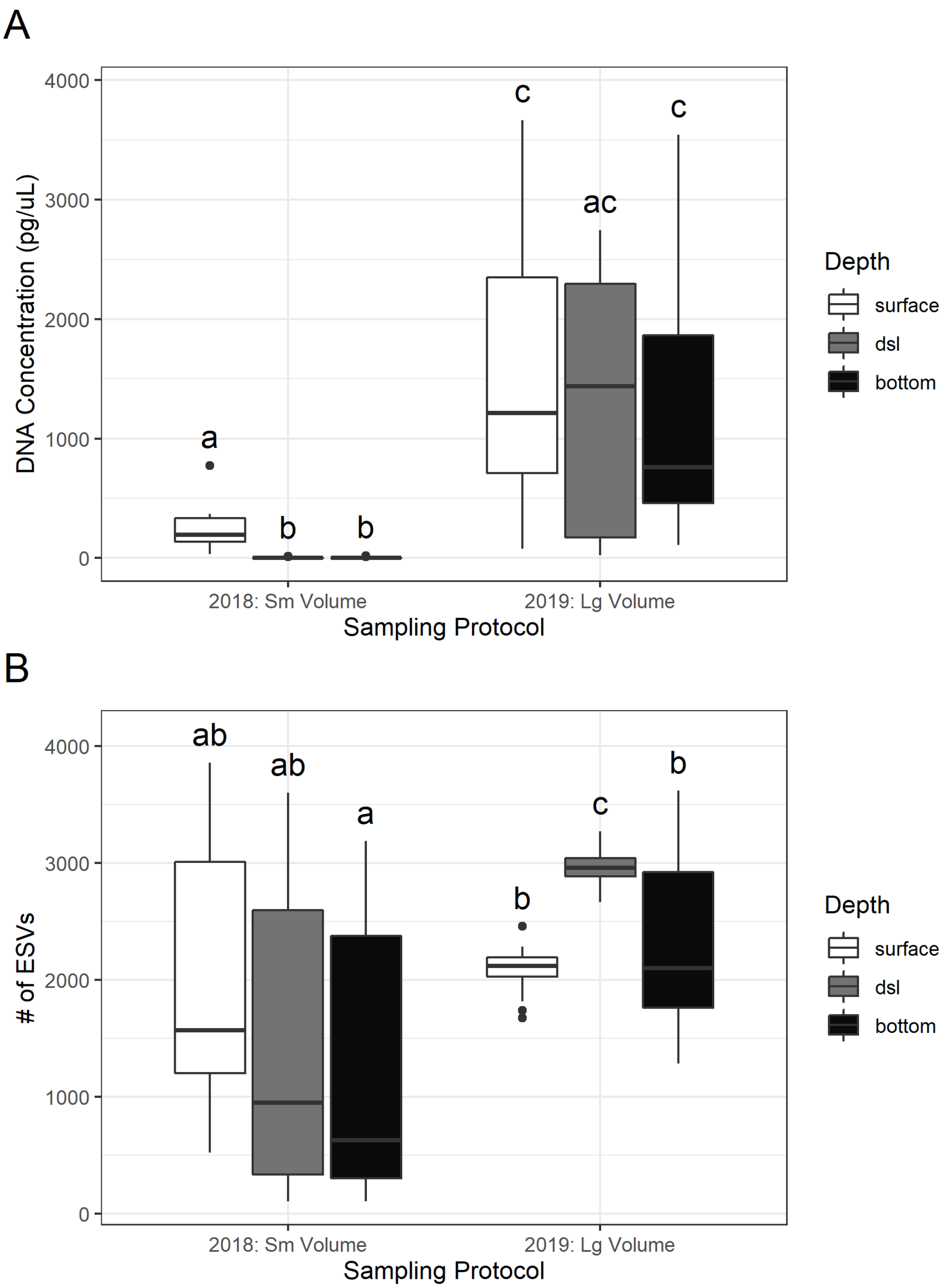


**S3 Fig. Comparison of (A) DNA concentration (pg/µL) in extracts and (B) number of ESVs recovered from small volume samples collected in 2018 and large volume samples collected in 2019 at various depth sampling locations (surface, deep scattering layer, bottom).** Different letters indicate significant differences.
